# *C. elegans* provide milk for their young

**DOI:** 10.1101/2020.11.15.380253

**Authors:** Carina C. Kern, StJohn Townsend, Antoine Salzmann, Nigel B. Rendell, Graham W. Taylor, Ruxandra M. Comisel, Lazaros C. Foukas, Jürg Bähler, David Gems

## Abstract

Adult *C. elegans* hermaphrodites exhibit severe senescent pathology that begins to develop within days of reaching sexual maturity (Ezcurra et al., 2018; Garigan et al., 2002; Herndon et al., 2002; Wang et al., 2018). For example, after depletion of self-sperm, intestinal biomass is converted into yolk leading to intestinal atrophy and yolk steatosis (pseudocoelomic lipoprotein pools, PLPs) (Ezcurra et al., 2018; Garigan et al., 2002; Herndon et al., 2002; Sornda et al., 2019). These senescent pathologies are promoted by insulin/IGF-1 signalling (IIS), which also shortens lifespan (Ezcurra et al., 2018; Kenyon, 2010). This pattern of rapid and severe pathology in organs linked to reproduction is reminiscent of semelparous organisms where massive reproductive effort leads to rapid death (reproductive death) as in Pacific salmon (Finch, 1990; Gems et al., 2020). Moreover, destructive conversion of somatic biomass to support reproduction is a hallmark of reproductive death (Gems et al., 2020). Yet arguing against the occurrence of reproductive death in *C. elegans* is the apparent futility of post-reproductive yolk production. Here we show that this effort is not futile, since post-reproductive mothers vent yolk through their vulva, which is consumed by progeny and supports their growth; thus vented yolk functions as a milk, and *C. elegans* mothers exhibit a form of lactation. Moreover, IIS promotes lactation, thereby effecting a costly process of resource transfer from postreproductive mothers to offspring. These results support the view that *C. elegans* hermaphrodites exhibit reproductive death involving a self-destructive process of lactation that is promoted by IIS. They also provide new insight into how the strongly life-shortening effects of IIS in *C. elegans* evolved.

---

*C. elegans* hermaphrodites are protandrous, producing first sperm and then oocytes, and reproduction ceases by around day 3 of adulthood due to self-sperm depletion. While working with adult hermaphrodites expressing vitellogenin (yolk protein) tagged with GFP (Grant and Hirsh, 1999), we noticed that older mothers leave patches of GFP-positive material on culture plates (Fig. 1a). Viewed under light microscopy these appeared as smears of a brownish substance (Fig. 1a). The presence of vented yolk protein was confirmed and found to be highest on days 4-6 of adulthood, immediately after cessation of egg laying, and then to continue at lower levels until at least day 14 (Fig. 1b; Extended Data Fig. 1a). Yolk was vented through the vulva in brief bursts, either alone or with unfertilised oocytes (Fig. 1c, Supplementary Video 1). Vented yolk also contains lipid, as shown by staining with a lipid dye (Fig. 1d; Extended Data Fig. 1b). Thus, cessation of egg laying due to self-sperm depletion is followed immediately by a burst of yolk venting.

**Fig. 1.**
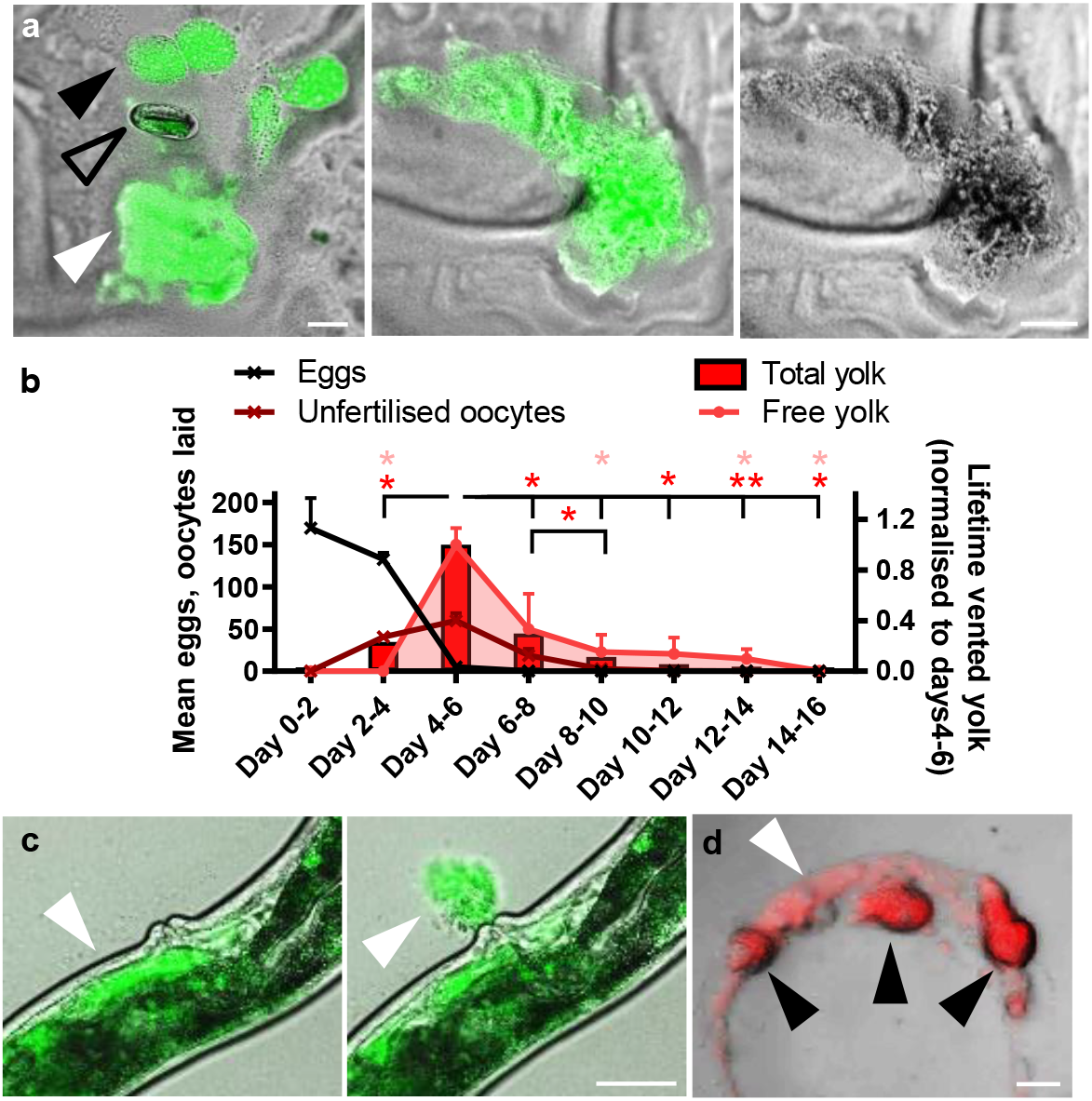
Post-reproductive *C. elegans* hermaphrodites vent yolk. **a**, Vented yolk pools and unfertilised oocytes on culture plates from hermaphrodites on d4 of adulthood, expressing *vit-2::GFP*. For other examples and separate images for Nomarski and epifluorescence microscopy see Extended data Fig. 1 c. **b**, Lifetime reproductive schedule, oocyte production, and proportion of total vented yolk (from oocytes + free yolk) and free yolk quantitated from VIT-2::GFP on plates and normalised to days 4-6. Mean ± s.e.m. of 3 trials displayed (n=50 worms per trial for venting and 10 per trial for brood sizes). \**P*<0.05, \*\**P*<0.01 by one-way ANOVA (Tukey correction; statistical tests performed on raw data), compared to day 4-6; red, total yolk (free + oocytes); pink, free yolk alone. **c**, Yolk vented through the vulva of a day 4 adult. Live imaging of VIT-2::GFP day 4 adults performed with yolk initially present in the uterus (left) and then seen vented 7 sec later (right) (white arrowhead). For presence of yolk in the uterus, other examples of yolk venting, time series, and comparison to egg laying on day 2 of adulthood see Extended data Fig. 1d-f and Supplementary video 1. **d**, Lipid in vented yolk and unfertilised oocytes from day 4 adults subjected to vital staining with the lipid dye Bodipy 493/503. Scale 50 μm. White arrowhead: yolk pools, black arrowhead: unfertilised oocytes, and open arrowhead: egg.

Given that yolk is a nutrient substance, one possibility is that vented yolk supports larval growth. Consistent with this, GFP-labelled yolk could be observed in the intestinal lumen of larvae (Fig. 2a). Notably, pre-treatment of *E. coli*-free agar plates with post-reproductive, 4-day old mothers enhanced growth of L1 larvae, relative to control plates (no pre-treatment, or pre-treatment with non-venting L3 larvae; Fig. 2b). Moreover, subjecting mothers to *vit-5,-6* RNAi, which prevents vitellogenin accumulation (Ezcurra et al., 2018) also blocked the benefit to wild-type larval growth of plate pre-conditioning with venting mothers (Fig. 2b). *vit-5* RNAi decreases levels of the YP170 vitellogenin species but RNAi of *vit-6* (which encodes YP115/YP88) increases them (Sornda et al., 2019). These treatments, respectively, suppressed and enhanced effects of pre-treatment with venting mothers on larval growth (Fig. 2b), implying that YP170 is the major nutrient source within vented yolk.

**Fig. 2.**
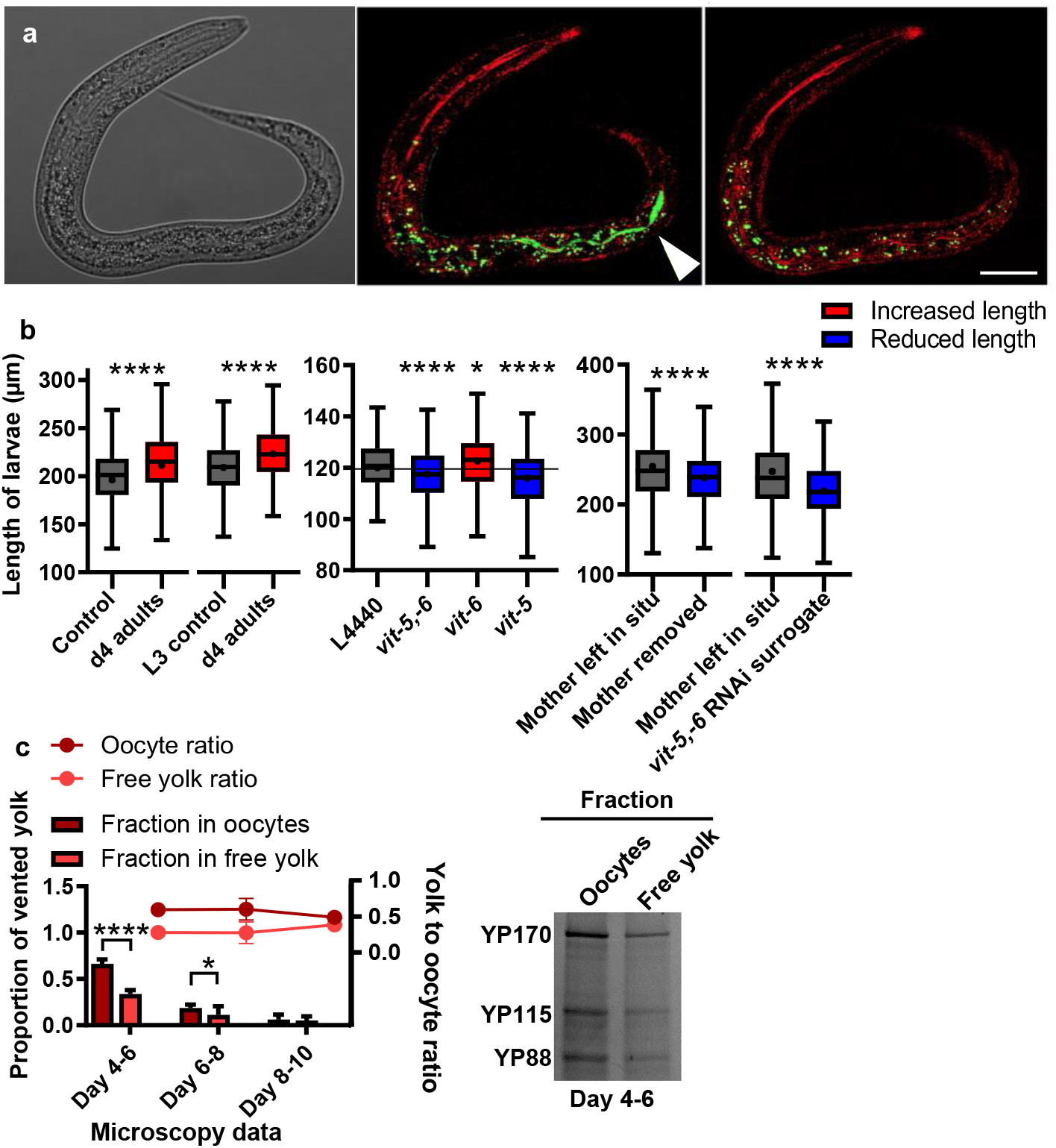
Vented yolk and unfertilised oocytes support larval growth. **a**, Yolk in the intestinal lumen of an L1 larva after being left on a plate with no food apart from vented yolk for 4 hr. Left: Nomarski microscopy image. Middle: larva imaged immediately after removal from plates using reflectance confocal microscopy (RCM) to highlight the refractive material of the terminal web that surrounds the intestinal lumen (fluorescence filters MBS T80/R20 and 405 nm excitation) (red) and superimposed airyscan image (488 nm excitation) (green) for GFP-labelled yolk. Right: same larva imaged 100 min later showing yolk no longer in the lumen, for comparison. White arrowhead: VIT-2::GFP. Scale bar 20 μm. Fluorescence outside intestinal lumen is from gut granule autofluorescence^30^. For details on RCM, see Extended Fig. 2b,c. **b**, Tukey box plots (line at median, + at mean) of length measurement of L1 larvae. Left and middle: larvae left from the egg stage for 48 hrs on plates preconditioned with day 4 adults left to vent for 24 hrs. L4440 empty vector control for RNAi-treated adults. Right: day 3 adults left to lay last few eggs and either left with larvae to vent yolk for 48 hrs or removed, or replaced with RNAi-treated surrogate mothers. Combined data of 3 trials (n=200 worms per trial). Kolmogorov–Smirnov non-parametric test. Red: treatments that increase the dependant variable; and blue: that reduce the dependant variable. All larvae in trials are wild type. **c**, Relative levels of free yolk vs yolk in unfertilised oocytes quantitated from VIT-2::GFP fluorescence on plates (n=10 worms per trial). Data normalised to total yolk on days 4-6. Mean ± s.d. of 3 trials displayed. Right: Protein gel showing YP bands after yolk collection from plates on d4-6. For protein gel data of vented free yolk on all days see Extended Data Fig. 2d. For chemotaxis data showing attraction towards adult hermaphrodites rather than vented yolk see Extended data Fig. 2e. \**P* <0.05, **** *P* <0.0001 by one-way ANOVA (Bonferroni correction).

Next we tested whether yolk vented by mothers can benefit their own larvae. Fully-fed 3 day old mothers were washed, placed on *E. coli*-free plates and left to lay their last eggs, and then either left *in situ* to vent yolk, or removed. Removal of mothers reduced growth of their progeny (Fig. 2b). In a further test, mothers were replaced after egg laying with other mothers of the same age but treated with *vit-5,-6* RNAi, and this abrogated the benefit to larval growth (Fig. 2b). Taken together, these findings show that after self-sperm depletion *C. elegans* mothers can enhance growth of their offspring by venting yolk through the vulva. As a means of transferring resources from mother to offspring after egg laying, vented yolk serves a function similar to that of mammalian milk; we therefore propose the term *yolk milk* to describe it. These findings reveal a surprising new function for the intestine and vulva in *C. elegans* hermaphrodites: that of lactation.

After sperm depletion hermaphrodites lay over 100 excess, unfertilised oocytes, the overall volume of which exceeds that of the hermaphrodite herself, which has been noted as oddly wasteful and futile (Ward and Carrel, 1979). The timing of unfertilised oocyte production is similar to that of yolk venting (Fig. 1b). Indeed, yolk and unfertilised oocytes are often vented together (Fig. 1a) suggesting a possible role for vented oocytes in yolk milk transport. Consistent with this, oocytes also contained large amounts of yolk, as shown by VIT-2::GFP, and confirmed by gel electrophoresis (Fig. 2c). At the outset of yolk/oocyte venting, oocytes contained twice as much vitellogenin as free vented yolk, but the proportion of the latter increased with age until day 10, when the ratio is 1:1 (Fig. 2c). To establish the extent to which vitellogenin delivered in each manner supports larval growth, free yolk and oocyte fractions were separated from conditioned plates on which L1 larvae had or had not been present for 24 hr, and vitellogenin content was assayed. The results show that larval feeding causes a reduction in yolk from both the oocyte and free yolk fractions (Extended Data Fig. 2a), i.e. vitellogenin in both free yolk pools and unfertilised oocytes was consumed by larvae. These results provide a possible explanation for the enigma of unfertilised oocyte production: that it represents an adaptation, aiding delivery of yolk milk to hungry young larvae (cf. milk and cookies).

*daf-2* insulin/IGF-1 receptor mutants have extended lifespan (Kenyon et al., 1993; Kimura et al., 1997), and show reductions in vitellogenin synthesis (Depina et al., 2011; Sornda et al., 2019), PLP accumulation (Ezcurra et al., 2018), and unfertilised oocyte production (Gems et al., 1998). Thus, IIS promotes ageing and production of yolk and oocytes. To test whether IIS promotes lactation we examined *daf-2* mutants, and found that yolk venting and promotion of wild-type larval growth by post-reproductive mothers was strongly reduced (Fig. 3a,b,c). Conversely, the *daf-2(gk390525)* gain-of-function (gf) mutation increased yolk venting and promotion of larval growth (Fig. 3a,b). Effects of *daf-2* on senescent pathology and lifespan require the *daf-16* FOXO transcription factor (Ezcurra et al., 2018; Kenyon et al., 1993). Accordingly, the *daf-16(mgDf50)* null mutation also restored yolk venting to *daf-2* mutants, and promotion of larval growth (Fig. 3a,b). Moreover, mutation of the *daf-18* PTEN phosphatase (Mihaylova et al., 1999), which increases phosphatidylinositol (3,4,5)-trisphosphate (PIP3) signalling, increased yolk venting and larval growth (Fig. 3a,b). *daf-2(e1370)* surrogate mothers also failed enhance growth of wild-type larvae hatched from final eggs (Fig. 3c; cf Fig. 2b right). In addition, suppression of unfertilised oocyte production by *daf-2(e1370)* is *daf-16* dependent (Gems et al., 1998), and numbers of unfertilised oocytes laid were increased by both *daf-2(gf)* and *daf-18(nr2037)* (Fig. 3d). These results imply that IIS promotes *C. elegans* lactation, through promotion of *vit* gene expression and of gut-to-yolk biomass conversion (Ezcurra et al., 2018). Thus, reduced IIS in *daf-2* mutants reduces later-life contributions to reproductive fitness and their associated costs, i.e. yolk milk production contributes to *C. elegans* senescence.

**Fig. 3.**
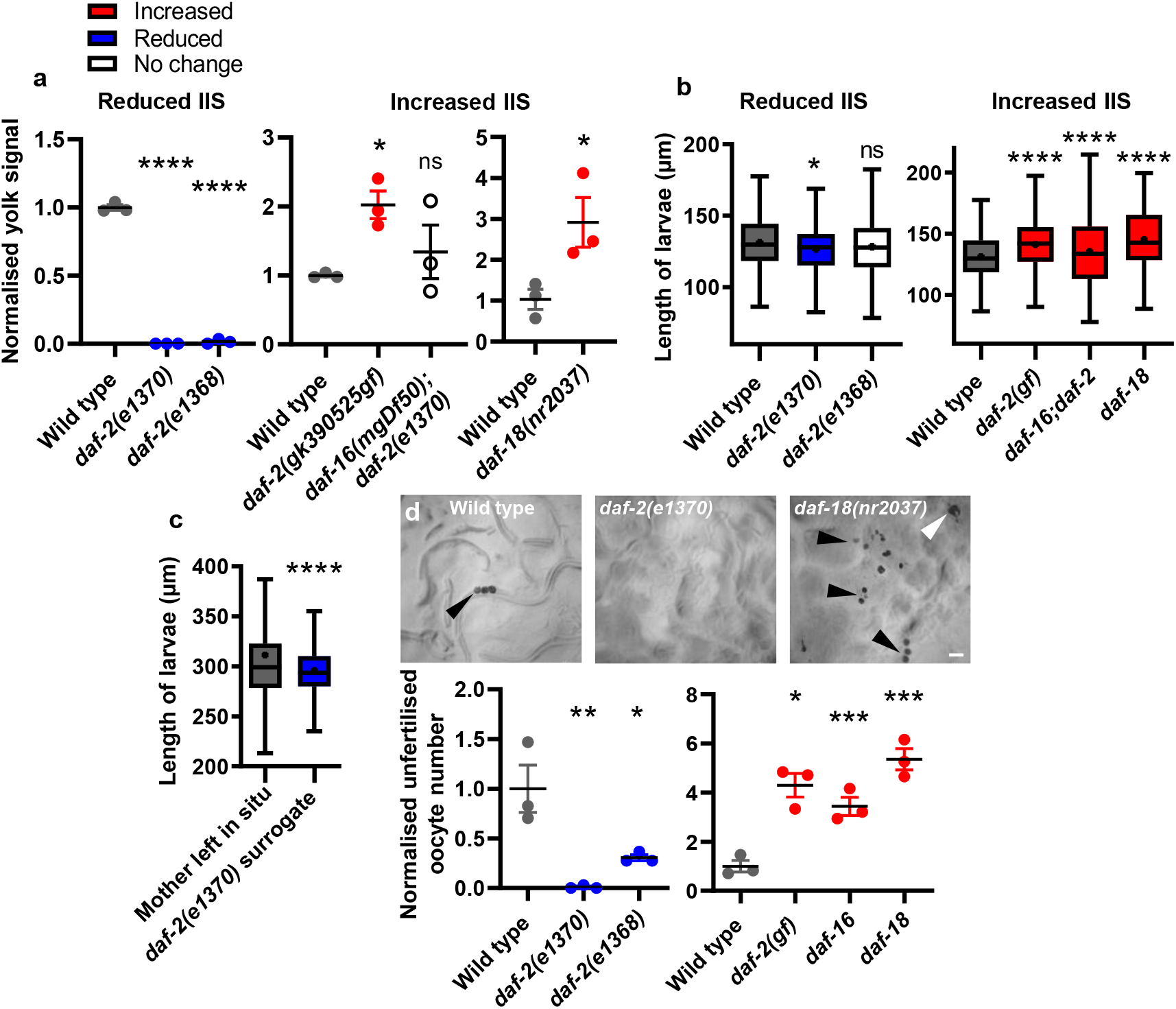
Yolk venting is promoted by insulin/IGF-1 signalling (IIS). **a**, Yolk venting is down-regulated by reduced IIS and up-regulated by enhanced IIS. Quantitated YP170 band on protein gels. **b**, Decreased yolk milk provisioning by reduced IIS reduces larval growth on preconditioned plates relative to wild-type, and vice versa for increased IIS. **c**, Decreased larval growth on plates with *daf-2(e1370)* surrogate mothers. Wild-type mothers were allowed to lay their last eggs and either left in situ or replaced. **d**, Unfertilised oocyte production is down-regulated by reduced IIS and up-regulated by increased IIS. Bottom: Number laid for 24 hrs on d4-6 of adulthood and normalised to wild type. Means ± s.e.m. of 3 trials (n=10 worms per trial). Top: NGM plates imaged after 15 wild-type, *daf-18(nr2037)* or *daf-2(e1370)* worms were left for 24 hrs. Black arrowheads: unfertilised oocytes; white arrowhead: yolk pool. Red: treatments that increase the dependant variable; blue: the reduce the dependant variable; white: no effect. \**P*<0.05, \*\**P*<0.01, \*\*\**P*<0.0001, \*\*\*\**P*<0.00001; one-way ANOVA (Dunnett’s correction) or unpaired t-test.

Our findings imply that the vitellogenin-rich fluid secreted by sperm-depleted *C. elegans* functions as a milk. To further characterise *C. elegans* yolk milk, secreted proteins were collected from L3 and 4 day old hermaphrodites, and subjected to proteomic analysis. The set of protein present in the latter but not the former represents an adult-specific secretome that includes abundant vulvally-vented proteins, and may also include proteins secreted via the excretory pore and anus, or from the nematode surface. A further expectation is that the secretome will include both proteins of functional significance (e.g. yolk milk proteins), and random proteins shed from internal organs. Our analysis defined a set of 125 secreted proteins. Given that vitellogenins are secreted proteins whose levels increase dramatically with age in an IIS-dependent fashion, we focused on proteins that were a) IIS regulated, b) more abundant with age, and c) secreted (possessing a predicted N-terminal signal peptide). Overall, 21 secretome proteins proved to be IIS regulated, 30 up-regulated in old age, and 28 to contain signal peptides (Fig. 4a,b, Extended data Table 1). Of these, 17 proteins exhibited all three features (Fig. 4c), which is 82-fold more than expected by chance alone, considering the frequency of each feature among all *C. elegans* proteins. Aside from the vitellogenins, which were the most abundant proteins, there were also seven transthyretin-related (*ttr*) proteins, TTR-2, −15, −16, −18, −30, −45 and −51. TTRs often function as carrier proteins for lipophilic compounds (Goodman, 1987; Woeber and Ingbar, 1968). Also present were FAR-3, another predicted lipid-binding protein, that is expressed in the vulva, and also FAR-1 and −2. Lipid carrier proteins are also abundant in mammalian milk, suggesting possible functional similarities. Thus, IIS promotes the production of certain proteins secreted from older worms whose abundance increases with age, defining a proposed yolk milk proteome.

**Fig. 4.**
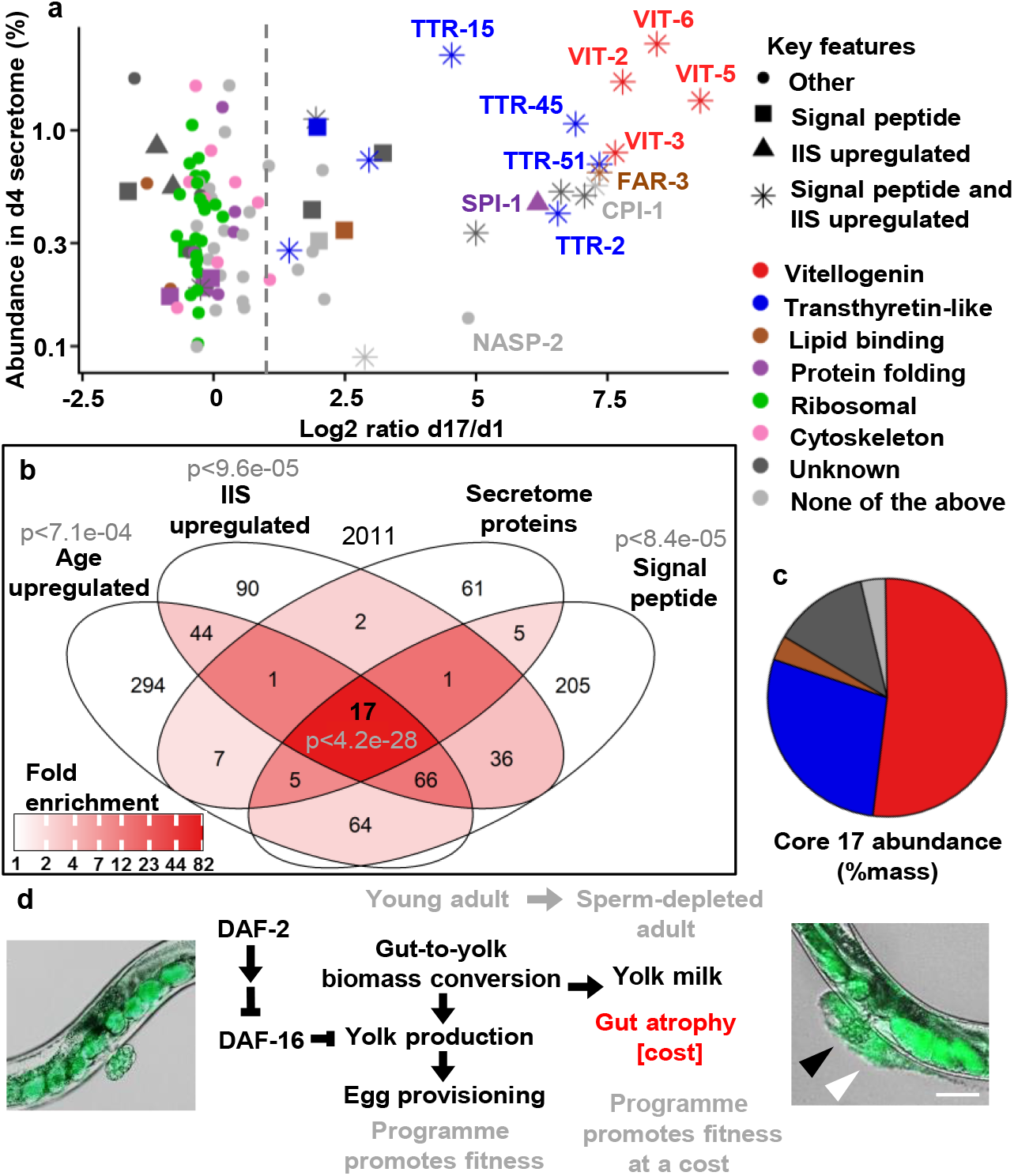
Proteomic analysis of adult secretome. **a**, Relationship between protein abundance in the day 4 adult secretome and increased abundance with age in the overall proteome, from published data for wild type and *daf-2(e1370)* (Walther et al., 2015). ‘Age upregulated’ was defined as showing an increase of >log2=1 day 17 vs day 1 (vertical dotted line). Right hand cluster includes many IIS- and age-upregulated secreted proteins. Left hand cluster likely includes proteins shed by tissue breakdown. **b**, Enrichment analysis of adult-specific secretome proteins relative to all proteins detected in the reference study (Walther et al., 2015). Significant over-representation was detected using a SuperExactTest (Wang et al., 2015). **c**, Percentage abundance categorisation of the core 17 proteins in the adult-specific secretome that are IIS upregulated, age upregulated and likely to be secreted (i.e. bear a predicted N-terminal signal peptide). For raw MS data see Supplementary File 1. For details of IIS upregulation and *daf-16* dependence see Extended Data Fig. 3. For L3 secretome and specific proteins of interest see Extended Data Fig. 4. For tissue enrichment analysis and comparison to proteomic analysis of human milk see Extended Data Fig. 5. **d**, Model of intestinal conversion to yolk in post sperm-depleted adults, where the benefit of yolk-milk comes at the cost of intestinal atrophy. Here IIS promotes self-destructive somatic biomass repurposing, thus promoting senescence in a fashion typical of organisms exhibiting semelparous reproductive death (Gems et al., 2020).

In this study, we show that post-reproductive *C. elegans* hermaphrodites exhibit a form of primitive lactation, releasing yolk milk (free and in oocytes) through their vulva that can support growth of progeny. We propose that this provide a fitness benefit, coupled to pathological changes to reproduction-associated organs, in a process of semelparous reproductive death (Fig. 4d). This conclusion is supported by comparative analysis of other *Caenorhabditis* species which shows that reproductive death is promoted by the germline in hermaphrodites but absent in unmated females (Kern et al., 2020), and by comparison with other organisms that exhibit reproductive death (Gems et al., 2020). We have argued elsewhere that reproductive death can facilitate the evolution of programmed adaptive death, which further shortens lifespan (Galimov and Gems, 2020; Lohr et al., 2019).

Milk feeding of larvae by mothers is a previously undescribed feature of *C. elegans* life history. Such maternal care by nematodes might seem surprising, but lactation occurs in other invertebrates, including tsetse flies (*Glossina* spp.) (Benoit et al., 2015) and the Pacific beetle cockroach *Diploptera punctata* (Marchal et al., 2013). Milk feeding by *C. elegans* also resembles trophallaxis, the transfer of food or nutritious fluids between individuals in a community, e.g. in social insects (LeBoeuf, 2017). It is likely that in the wild, reproducing *C. elegans* exist largely as colony-like, high density, clonal populations (Lohr et al., 2019; Schulenburg and Félix, 2017). Potentially, it is into this collective that sperm-depleted hermaphrodites secrete yolk milk.

Hermaphroditism facilitates rapid colonization of new food patches (Schulenburg and Félix, 2017), but protandry leaves mothers unable to contribute to fitness after sperm depletion, an example of a *bauplan*-type trade off (Gould and Lewontin, 1979; Lohr et al., 2019). We suggest that later-life yolk milk production is an adaptation to circumvent this block to continued maternal contribution to fitness, here inclusive fitness of the surrounding clonal population. Lactation also provides a solution to the long-standing enigma of copious unfertilised oocyte production (Ward and Carrel, 1979).

Ageing is thought to evolve partly because genes exhibit antagonistic pleiotropy (AP), exerting both beneficial and deleterious effects on fitness; if the latter only occur later in life, such AP genes may be favoured by natural selection (Williams, 1957). Previously, post-reproductive yolk production and the pathologies to which it is coupled have been interpreted as futile run-on of reproductive function (Ezcurra et al., 2018; Herndon et al., 2002; Sornda et al., 2019). This is consistent with the AP theory, and recent ideas about its proximate mechanisms in programmatic terms, which argue that futile run-on of biological programmes in later life (or quasi-programmes) contributes to senescent pathology (Blagosklonny, 2006; de Magalhães and Church, 2005; Williams, 1957). But if late-life yolk production contributes to fitness, then such production is not a quasi-programme but a programme proper, and pathologies such as gut atrophy are a direct cost of reproduction (Speakman, 2008). Thus, there is a trade-off between increased fitness due to yolk milk feeding of larvae (benefit) and intestinal atrophy leading to adult mortality (cost) (Fig. 4d). These results provide a fresh account of the AP action of IIS pathway genes such as *daf-2*.

## Methods

See supplementary information.

## Supporting information

Supplementary Guide

Supplementary Folder 1

## Acknowledgments

We thank L. Cao, Y. de la Guardia, N. Hui, S. Ranasinghe, N. Sergent, M. Cuffaro and N. Thant for technical assistance and minor contributions, and N. Alic, J. Labbadia, S.-J. Lee, E. Murphy, T. Niccoli and S. Sumner for useful discussion, and/or comments on the manuscript. Some strains were provided by the Caenorhabditis Genetics Center, which is funded by NIH Office of Research Infrastructure Programs (P40 OD010440). S.T. was supported by a Boehringer Ingelheim Fonds PhD Fellowship. Core support for the Wolfson Drug Discovery Unit is provided by the UK National Institute for Health Research Biomedical Research Centre and Unit Funding scheme via the UCLH/UCL Biomedical Research Centre. Funding for the proteomics platform was provided by the Wolfson Foundation and the UCL Amyloidosis Research Fund. This work was supported by a Wellcome Trust Strategic Award (098565/Z/12/Z) and a Wellcome Trust Investigator Award (215574/Z/19/Z) to D.G.

## Author contributions

D.G. supervised the project. C.C.K. and D.G. conceived the project, designed the experiments, and wrote the manuscript. A.S., R.M.C., L.F. and C.C.K. performed behavioral experiments. N.R. and G.T. performed MS analysis. D.G., C.C.K. and S.T. designed and performed MS data analysis. J.B. assisted in the interpretation of MS data. J.B. and L.F. contributed to editing the manuscript.

