## Extended Data and Methods for "*C. elegans* provide milk for their young"

### Supplementary Information Guide

<u>**This file**</u>

**Methods.**

**Extended Data Figs. 1-5.**

**Extended Table 1.**

<u>**Additional files**</u>

**Supplementary Video 1 | Representative video of yolk venting by d4 hermaphrodite.** A pool of yolk, marked with VIT-2::GFP, is initially visible in the uterus near the vulva. There follow several very small bursts of venting and then one large burst, accompanied by uterine muscle contractions, after which the uterine yolk pool is depleted.

**Supplementary File 1 | d4 adult-specific and L3 secretome.** Mass spectrometry raw data for proteins secreted into the surrounding medium by *C. elegans*.

**Supplementary Folder 1 | Enrichment of proteins with Interpro and GO terms in d4 adult-specific secretome and human milk proteome.**

#### Methods

No statistical methods were used to predetermine sample size. The experiments were not randomised. The investigators were not blinded to allocation during experiments and outcome assessment.

##### Culture methods and strains

*C. elegans* maintenance was performed using standard protocols (Brenner, 1974). Unless otherwise stated, all strains were grown at 20˚C on nematode growth media (NGM) with plates seeded with *E. coli* OP50 to provide a food source. An N2 hermaphrodite stock recently obtained from the Caenorhabditis Genetics Center was used as wild type (N2H) (Zhao et al., 2019). Genotypes of mutants used are as described in Wormbase (www.wormbase.org). Strains used included DR1296 *daf-2(e1368)*, GA114 *daf-16(mgDf50)*; *daf-2(e1370)*, GA1500 *bIs1 [pvit-2::vit-2::GFP + rol-6(su1006)]*, GA1928 *daf-2(e1370)*, GR1307 *daf-16(mgDf50)*, NS3227 *daf-18(nr2037),* RT130 *pwIs23 [vit-2::GFP]*, and VC20747 *daf-2(gk390525*).

##### Live imaging and video capture of venting behaviour

In order to impede worm locomotion and facilitate imaging, polybead microspheres were used. d4 RT130 adults were placed on an NGM plate (no bacteria) in a 15-20 µl drop of 0.1 µm non-fluorescent polybead microspheres (2.5% solids [w/v] aqueous suspension with a coefficient of variance of 15%, 4.55 x $10^{13}$ particles/ml [Polysciences]). To remove any aggregated particles these were previously filtered through a 0.5 µm pore size syringe filter. A coverslip was then gently placed over the animal and observations made using a 20x objective (200x magnification) and Nomarski optics. To prevent fluorescence bleaching, Nomarski/GFP superimposed videoing and imaging were commenced only when movement of the vulval muscles was observed, which often preceded yolk milk venting or laying of eggs or unfertilised oocytes.

##### Nomarski and epifluorescence microscopy imaging

Unless otherwise stated, live worms were placed onto 2% agar pads and anaesthetised in a drop of 0.2% levamisole, with coverslips gently placed on top. Images were captured using either a Zeiss Axioskop 2 plus microscope with a Hamamatsu ORCA-ER digital camera C4742-95 and Volocity 6.3 software (Macintosh version) for image acquisition; or an ApoTome.2 Zeiss microscope with a Hamamatsu digital camera C13440 ORCA-Flash4.0 V3 and Zen software. A constant exposure time was maintained between samples in fluorescence intensity comparisons. Brightness and contrast were adjusted equally across the entire image, and where applicable applied equally to controls. Where Nomarski and fluorescence images were superimposed, brightness and contrast were adjusted separately prior to superimposition.

##### Confocal and airyscan imaging

For this an inverted LSM880 microscope equipped with an Airyscan detector (Carl Zeiss, Jena) was used with a Plan-Aprochromat 63 x 1.4 [numerical aperture (NA)] oil objective with a working distance of 0.19 mm. A 488 nm Argon laser was used for GFP excitation. In confocal mode, emission was recorded with an inbuilt GaAsP detector. For airyscan, the emission was recorded with the in-built 32-element GaAsP detector. For images showing sample change over time, images were only taken around every 50 min to prevent sample photobleaching. In the acquisition of 3D z-series, samples were imaged up to a sample depth of 41 μm, with images of 41 z-planes taken evenly through half of the diameter (dorsoventral) of the nematode. Data was processed using Fiji software (NIH), and the 3D Viewer plugin was used for 3D reconstruction. Brightness and contrast were adjusted equally across the entire image, and where applicable applied equally to controls. Where Nomarski and fluorescence images were superimposed, brightness and contrast were adjusted separately prior to superimposition.

**Reflectance confocal microscopy**

This was performed using an inverted LSM880 microscope (Carl Zeiss, Jena) and a Plan-Aprochromat 63x 1.4 numerical aperture (NA) oil objective with a working distance of 0.19 mm. The main beam splitter was set to T80/R20 with multiphoton laser 405 nm excitation. Emission was recorded using an inbuilt GaAsP detector.

**Fluorescence quantitation of vented yolk milk**

10 GA1500 L4 larvae were placed on 35 mm NGM plates (n = 5 plates per trial; total 50 worms) seeded with 100 μl *E. coli* OP50 and transferred every 24 hr to new plates. After transferring, 5 superimposed GFP/Nomarski 50x magnification images were taken of each plate at random positions across the bacterial lawn to sample approximately half of the lawn area. For each time point 3 control NGM plates (no worms) were treated in the same way and imaged.

Fluorescence quantitation of images was performed using the formulae below to account for varying levels of NGM background fluorescence in different plates and variations in fluorescence excitation. Yolk milk pools form localised regions of bright fluorescence on plates. Therefore the minimum emission fluorescence of an NGM plate can be used as an indicator of the true level of background fluorescence, since the range between the minimum and maximum background fluorescence of NGM is relatively constant between plates (data not shown). By measuring the background autofluorescence of control plates and normalising it to the minimum of treated plates, an accurate estimate of the range and maximum background fluorescence of treated control plate can be estimated and subtracted, using the following procedure (note: fluorescence calculations only refer to the GFP channel).

(i) Images with dust and/or cholesterol crystals (which can reduce the minimum emission fluorescence and affect the background fluorescence range) were manually censored (visualised under Nomarski as black dots or black crystals).

(ii) The ratio of minimum and maximum values of fluorescence intensity (FIR) is given by:

$$\rho = \frac{F_{\max}}{F_{\min}}$$

where $F_{\max}$ is the maximum emission fluorescence detected on a plate, and $F_{\min}$ the minimum emission fluorescence detected (both from background fluorescence; $F_{\min}$ to $F_{\max}$ = background fluorescence range).

(iii) The FIRs were averaged for the three control plates imaged each day to provide a Daily Control Fluorescence Intensity Ratio $\bar{\rho}$ (DCIR). This also accounts for variation in UV lamp output during sample excitation. For each treated plate image, the DCIR was multiplied by the image $F_{\min}$ in order to set a threshold (T) fluorescence level on treated plates below which all fluorescence was considered background fluorescence and ignored.

$$F_T = F_{\min}\bar{\rho}$$

(iv) Once done, the remaining fluorescence from above the threshold was isolated to ROIs containing GFP-labelled protein, with the fluorescence consisting of both GFP fluorescence as well as background fluorescence. Background fluorescence from these ROIs was removed follows:

$$F_{\mathrm{GFP}} = -AF_T + \sum_u F_u \qquad A = \text{Pixel count}$$

where $F_{(\mathrm{GFP})}$ is the total fluorescence from GFP on the plate, and $F_u$ is the fluorescence from a given ROI, with $\sum_u F_u$ the total fluorescence from the collection of ROIs remaining on an image after step (iii).

(v) Separation of yolk and oocyte fluorescence was done manually for each image using Volocity 6.3 Acquisition. The free-drawing tool was used to identify oocytes on Nomarski/GFP images by viewing the Nomarski image, followed by subtracting all sum fluorescence within the ROIs from the total calculated in steps (i-iii).

$$F_{\mathrm{Free\ yolk}} = F_{\mathrm{GFP}} - \sum_u F_{\mathrm{Oocyte}(u)}$$

**Gel electrophoresis and quantitation of vented yolk proteins**

100-200 L4 larvae were maintained on 35 mm plates freshly seeded with 100 µl OP50. Following transfer each 24 hr, yolk milk was washed off with 1 ml of M9 containing 0.001% NP-40 to solubilise vitellogenin (Sharrock, 1983) and 2 µg/ml BSA as an external standard. This solution was pipetted onto plates and a polystyrene bacterial loop was then used to suspend all surface material, including the bacterial lawn, in the solution. The resulting suspension was pipetting out and into an Eppendorf tube. Inclusion of BSA served a dual purpose as a control for protein loss during sample collection from plates, and as a loading control for gel electrophoresis. Samples were spun at 2,600 rpm for 10 min at 4˚C, and the pellet collected as the oocyte-containing fraction which contained the majority of the *E. coli*, and 500 µl of the supernatant for free yolk milk analysis. Bacterial protein bands did not interfere with the detection of YP170 or BSA bands analysed (Extended Data Fig. 1a). Samples were then lyophilised with 10 µl glycerol, and the remaining pellet solubilised in a solution of 30 µl 4% SDS solution pH9, 1 ml 1M Tris HCL pH 8, 5 ml Milli-Q water, 10 µl 0.5M EDTA, and 12 mg bromophenol blue (method optimised for YPs) by heating 2-4 times (based on the presence of a pellet post treatment) at 95˚C for 5 min while vortexing periodically, and finally centrifuged at 6,000 rpm for 15 min. To prevent sample loss, all pipetting was performed using LoBind pipette tips and storage was in LoBind Eppendorf tubes. Sodium dodecyl sulfate–polyacrylamide gel electrophoresis (SDS-PAGE) was then performed, using Criterion XT Precast Gels 4–12% Bis-Tris (Invitrogen) and XT MOPS (Invitrogen) as a running buffer (7:1 ratio with Milli-Q water) at 90 V. Gels were stained with colloidal Coomassie blue as described (Kang et al., 2002), using 5% aluminium sulphate-(14-18)-hydrate and 2% orthophosphoric acid (85%) to create colloidal particles. Gels were analyzed using ImageQuant LAS 4000 (GE Healthcare). Protein band identification was based on published data (Depina et al., 2011). Within lanes, vented YPs were normalised to the BSA that had been added during sample collection as an external standard.

**Staining of lipid in hermaphrodites prior to venting**

For each time point, vital staining of worms was performed as described (Klapper et al., 2011) using the fluorescent dye BODIPY 493/503 (488 nm Exc./505-575 nm Em.) (Invitrogen) at a final concentration of 6.7 µg/ml, with hermaphrodites incubated in dye for 20 min at RT in darkness. Nematodes were then transferred to NGM plates (no *E. coli*) and allowed to crawl for 3-5 min to remove surface dye, followed by transfer to 35 mm NGM plates (no *E. coli*), with 20 worms per plate (1-2 plates per trial), and left to vent for 24 hr. Fluorescence quantitation was performed as described for fluorescence quantitation of vented YPs, with the exception that Nomarski/fluorescence superimposed images were taken of the 5 regions with the most BODIPY fluorescence. This was due to the lower levels of BODIPY fluorescence relative to VIT-2::GFP fluorescence (likely due to differences in the fluorophores rather than the quantity of vented lipid vs protein). Control plates with L4 larvae (rather than adults) contained no patches of BODIPY fluorescence, ruling out a contribution of excess surface dye or egested dye to observed BODIPY staining.

**Determination of brood size and unfertilised oocyte number**

L4 hermaphrodites were placed individually on 35 mm NGM plates with OP50 (n = 10 worms per trial), and worms transferred every 24 hr until unfertilised oocyte production ceased. Eggs were left for 24 hr to hatch and brood sizes counted. Clumps of unfertilised oocytes were carefully separated using a platinum pick to distinguish individual cells, using a dissecting microscope at maximum magnification.

**Chemotaxis assay**

The chemotaxis assay was adapted from a previous study (Margie et al., 2013). Briefly, a 5 cm NGM plate was divided into quadrants with test compounds and d5 adults placed on two opposite quadrants and controls placed on the 2 remaining quadrants. Plates were then left for 4 hr to form a chemotactic gradient. All spots also contained 4 µl of 0.25 M sodium azide as anaesthetic. Next a 2 µl drop containing ~75 arrested L1 larva/µl was pipetted onto the centre of the plate within a 0.5 cm radius inner circle. Based on larval movement, a chemotaxis index (CI) was calculated using a formula as previously described (Margie et al., 2013).

For experiments involving adult worms, a single, heat-killed d5 adult was used for the treatment with controls treated the same way but with no worm. For experiments involving vented yolk, sample collection was performed by concentration of 100 d4 adults (washed twice to remove surface bacteria) onto a small microwell cell culture plate (Thermo Scientific Nunc 24-well cell-culture multidishes, 1.9 $cm^2$) containing 500 µl NGM lacking bactopeptone (to make the agar harder and limit bacterial growth) and carbenicillin added topically 24 hr prior to a concentration of 50 mM. This was done by picking worms in groups of 30-40 and holding them above the microwell, immediately after which a 10 µl drop of M9 was pipetted onto the pick to wash the worms into the well (no more than 30 µl of M9 was used per well). The worms were then allowed to vent for 24 hr, after which a sterile scalpel was used to cut around the NGM well and transfer it to a 10 ml NGM plate right side up. Adults were left for 1 hr to crawl off, after which remaining adults were picked off. The NGM with vented yolk was then cut into quadrants and each was placed as a treated section in a chemotaxis assay. Control sections were treated the same way but either had no worms, or L3 larvae added instead of d4 adults.

**Preconditioning of plates with vented yolk milk for larval growth assays**

30 d4 adults were washed twice in M9 to remove surface bacteria and left to vent for 24 hr on 35 mm NGM plates lacking bactopeptone, with 30 µl of 500 mM carbenicillin in Milli-Q water (plus 4 µl of 500 mM kanamycin in Milli-Q water for RNAi-treated adults as RNAi clones contain a carbenicillin/ampicillin resistance plasmid) added topically 24 hr prior (2-3 plates per trial). Controls had no d4 adults or L3 larvae treated in the same way. Plates with bagging (i.e. with internally hatched larvae) or ruptured worms were censored. Next, 200 alkaline hypochlorite-treated eggs were placed on each plate and left for 48 hr to develop, after which larval length was measured.

**Tests of yolk milk feeding by mothers of their own larvae**

Shortly after reaching d3 of adulthood, 30 hermaphrodites were washed twice in M9 and placed on 35 mm bactopeptone-less NGM plates (2-3 plates per trial) and left for 24 hr to lay their last eggs. Following this, the adults were either left in situ to vent yolk, removed, or replaced with surrogate mother worms of the same age. After 48 hr all adults were removed and larval length measured. Plates with bagging or ruptured worms were censored.

**Larval length measurements**

To collect larvae, plates were washed using 500 µl of M9 x 3 times and the collected liquid freeze-thawed (-80˚C) to straighten larvae. The samples centrifuged down at 2,000 RPM at 4˚C for 5 min and the pellet along with 500 µl of liquid above it placed on slides for imaging. Volocity 6.3 software (Macintosh version) was then used to measure length.

**Proteomic analysis of d4 hermaphrodite secretome**

350 fully-fed d4 hermaphrodites were picked onto NGM plates (no *E. coli*) and allowed to crawl for 30-60 sec to remove surface bacteria before being transferred to a 500 µl solution of 0.001% NP-40 in M9 to prevent vented yolk from adhering to the nematode surface (Sharrock, 1983). The worms were allowed to vent for 30 min (with picking of worms into each tube taking an additional 30 min) with the Eppendorf tubes maintained horizontally to aid diffusion of secreted proteins. The Eppendorf tube was placed upright for 1 min to allow nematodes to settle to the bottom of the tube, and 350 µl of solution drawn from the top of the tube. After collection, samples were carefully checked for the presence of adult worms and unfertilised oocytes, but none were observed. The adults worms were checked for the presence of ruptured animals, which were easy to identify, where found the sample collected from that tube was discarded. Secreted proteins were also collected for L3 larvae, as a negative control for venting through the vulva (which is absent at this stage).

Independent samples were digested with trypsin and prepared for proteomics analysis (Canetti et al., 2020). Samples were analysed on a Thermo Scientific Q-Exactive Plus Orbitrap mass spectrometer connected to an Ultimate 3000 nanoLC system. Samples were trapped on a Thermo Scientific Acclaim PepMap C18 cartridge (0.3 mm x 5 mm, 5 µm/100 Å) and then chromatographed on a Thermo Scientific Easy-Spray Acclaim PepMap C18 column (75 µm x 15 cm, 3 µm/100Å packing) eluting at 300 nl/min with a 30 min linear gradient of acetonitrile:water:formic acid (5:95:0.1 – 56:44:1 v/v/v). A full MS scan (m/z 135 – 2000 at 70,000 resolution) was acquired with a maximum injection time of 100 ms, and the 10 most intense ions with an intensity threshold 2.0e4 are selected for higher-energy C-trap dissociation

(HCD) with a lock mass of m/z 445.12003. The normalised collision energy is 30, with an isolation width of 2 Da and dynamic exclusion of 20 s; singly charged ions were excluded. All chromatography solvents were Optima LCMS grade (Fisher Scientific).

**Bioinformatic analysis of the d4 hermaphrodite secretome**

Proteomes were quantified using MaxQuant 1.6.12.0 (Cox and Mann, 2008) with the default search settings and the *C. elegans* protein database from Uniprot (downloaded 20 December 2019), and downstream analysis was performed using the Proteus package in R (Gierlinski et al., 2018). Proteins were considered detected in a sample if at least two proteotypic peptides from that protein were detected. Proteins were scored as present in the secretome if present in at least two of the three replicate samples. Proteins were only considered specific to the day 4 secretome if not detected in all three L3 control samples.

To estimate the composition of the secretome, proteins were manually classified according to known functions, and the sum of all peptide intensities across each grouping was used to calculate an approximate relative abundance of each grouping. Significant over-representation of secretome proteins with respect to other datasets was detected using a SuperExactTest (Wang et al., 2015). To assesses whether the adult secretome showed any distinct patterns of expression in ageing worms or IIS knockdown (in *daf-2(e1370)* and *daf-16(mu86)* mutants, which are similar to *daf-16(mgDf50)* mutants used in other experiments), an existing proteome dataset (Walther et al., 2015) was used. Proteins were considered upregulated in ageing worms if expression at least doubled from 1 day old worms to 17 day old worms. Proteins were considered IIS-upregulated if expression in 17 day old *wild-type* worms was at least twice the expression in 17 day old *daf-2(e1370)* worms, normalising for any initial expression differences between 1 day old wild-type and *daf-2(e1370)* worms. The presence of signal peptides in *C. elegans* proteins was predicted using SignalP 5.0 (Almagro Armenteros et al., 2019). In order to compare the composition of secretome proteins to human milk (D'Alessandro et al., 2010), cross-species gene set analysis was performed using the XGSA package in R (Djordjevic et al., 2016), accounting for protein homology mapping between *C. elegans* and humans. Tissue enrichment analysis was performed using the Wormbase tissue enrichment tool (Angeles-Albores et al., 2016) whilst GO term and Interpro term enrichment was performed using DAVID 6.8 (Huang et al., 2009).


Almagro Armenteros, J.J., Tsirigos, K.D., Sønderby, C.K., Nordahl Petersen, T., Winther, O., Brunak, S., von Heijne, G., Nielsen, H., 2019. SignalP 5.0 improves signal peptide predictions using deep neural networks. Nat. Biotech. 37, 420–423.

Angeles-Albores, D., Lee, R., Chan, J., Sternberg, P., 2016. Tissue enrichment analysis for *C. elegans* genomics. BMC Bioinformatics 17, 366.

Brenner, S., 1974. The genetics of *Caenorhabditis elegans*. Genetics 77, 71-94.

Canetti, D., Rendell, N.B., Gilbertson, J.A., Botcher, N., Nocerino, P., Blanco, A., Di Vagno, L., Rowczenio, D., Verona, G., Mangione, P.P., Bellotti, V., Hawkins, P.N., Gillmore, J.D., Taylor, G.W., 2020. Diagnostic amyloid proteomics: experience of the UK National Amyloidosis Centre. Clin. Chem. Lab. Med. 58, 948-957.

Cox, J., Mann, M., 2008. MaxQuant enables high peptide identification rates, individualized p.p.b.-range mass accuracies and proteome-wide protein quantification. Nat. Biotech. 26, 1367-1372.

D'Alessandro, A., Scaloni, A., Zolla, L., 2010. Human milk proteins: an interactomics and updated functional overview. J. Proteome Res. 9, 3339-3373.

Depina, A., Iser, W., Park, S., Maudsley, S., Wilson, M., Wolkow, C., 2011. Regulation of *Caenorhabditis elegans* vitellogenesis by DAF-2/IIS through separable transcriptional and posttranscriptional mechanisms. BMC Physiol. 11, 11.

Djordjevic, D., Kusumi, K., Ho, J., 2016. XGSA: A statistical method for cross-species gene set analysis. Bioinformatics 32, i620-i628.

Gierlinski, M., Gastaldello, F., Cole, C., Barton, G., 2018. Proteus: an R package for downstream analysis of MaxQuant output. BioRxiv, 416511.

Huang, D., Sherman, B., Lempicki, R., 2009. Systematic and integrative analysis of large gene lists using DAVID bioinformatics resources. Nat. Protoc. 4, 44-57.

Kang, D., Gho, S.G., Suh, M., Kang, C., 2002. Highly sensitive and fast protein detection with coomassie brilliant blue in sodium dodecyl sulfate-polyacrylamide gel electrophoresis. Bull. Korean Chem. Soc. 11, 1511-1512.

Klapper, M., Ehmke, M., Palgunow, D., Bohme, M., Matthaus, C., Bergner, G., Dietzek, B., Popp, J., Doring, F., 2011. Fluorescence-based fixative and vital staining of lipid droplets in *Caenorhabditis elegans* reveal fat stores using microscopy and flow cytometry approaches. J. Lipid Res. 52, 1281-1293.

Margie, O., Palmer, C., Chin-Sang, I., 2013. C. elegans chemotaxis assay. J Vis Exp, e50069.

McGhee, J.D., 2007. The *C. elegans* intestine, in: The *C. elegans* Research Community (Ed.), WormBook.

Sharrock, W., 1983. Yolk proteins of *Caenorhabditis elegans*. Dev. Biol. 96, 182-188.

Spieth, J., Blumenthal, T., 1985. The *Caenorhabditis elegans* vitellogenin gene family includes a gene encoding a distantly related protein. Mol. Cell. Biol. 5, 2495-2501.

Walther, D., Kasturi, P., Zheng, M., Pinkert, S., Vecchi, G., Ciryam, P., Morimoto, R., Dobson, C., Vendruscolo, M., Mann, M., Hartl, F., 2015. Widespread proteome remodeling and aggregation in aging *C. elegans*. Cell 161, 919-932.

Wang, M., Zhao, Y., Zhang, B., 2015. Efficient test and visualization of multi-set intersections. Sci. Rep. 5, 16923.

Zhao, Y., Wang, H., Poole, R.J., Gems, D., 2019. A *fln-2* mutation affects lethal pathology and lifespan in *C. elegans*. Nat. Commun. 10, 5087.

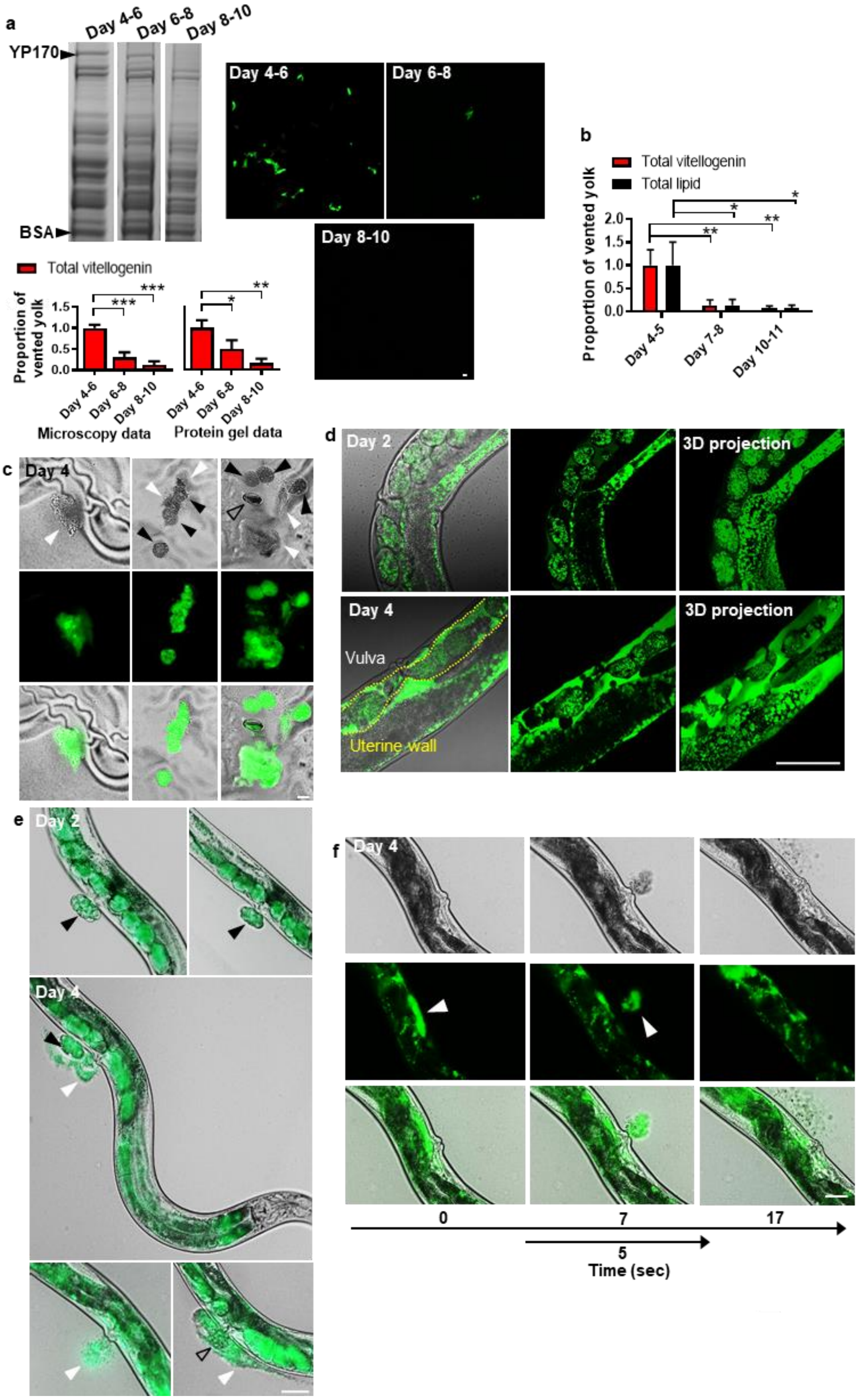


**Extended Data Fig. 1 | Characterising yolk venting. a**, Vented yolk on different days of adulthood. Left: Coomassie stained gel showing vented proteins (n=100 worms per trial); YP170 band is visible. BSA, external standard. Other YP bands are too faint to be seen. Right: yolk patches left on plates by VIT-2::GFP d4 adults left for 24 hr (n=10 worms per trial). Bottom: Relative levels of total yolk quantitated from VIT-2::GFP fluorescence on plate, or from protein gel analysis of YP170. Data normalised to total yolk on days 4-6. Mean ± s.e.m of 3 trials displayed. **b**, Lipid venting by BODIPY 493/503 vital stained adults follows the same pattern through time as vitellogenin venting by VIT-2::GFP adults. Values show mean ± s.e.m. of 3 trials. $P$ *<0.05, $P$ **<0.01 by one-way ANOVA (Bonferroni correction). **c**, Nomarski, GFP and superimposed images of vented yolk pools and unfertilised oocytes on NGM plates. **d**, Yolk in the uterus of d4 adults but little/negligible yolk in the uterus of d2 adults. VIT-2::GFP animals. Left: internal yolk airyscan confocal and Nomarski superimposed images; middle: airyscan confocal image through mid-body of worm; and right: 3D projection of images of 41 Z-planes taken up to a sample depth of 41 µm through half the width of each worm. **e**, Yolk venting on d4 compared with negligible yolk surrounding eggs laid on d2 of adulthood. Yolk venting on d4 may also be accompanied with the worm laying its last few fertilised eggs, or laying of unfertilised oocytes. **f**, Nomarski, GFP and superimposed images of yolk venting through time. One venting burst takes approximately 5 sec; c.f. Supplementary Video 1. Left: yolk seen in the uterus at time 0 sec, middle: yolk venting at time 7 sec, and right: less yolk in the uterus post venting at time 17 sec. Scale 50 µm. White arrowhead: yolk pools, black arrowhead: unfertilised oocytes, and open arrowhead: eggs.

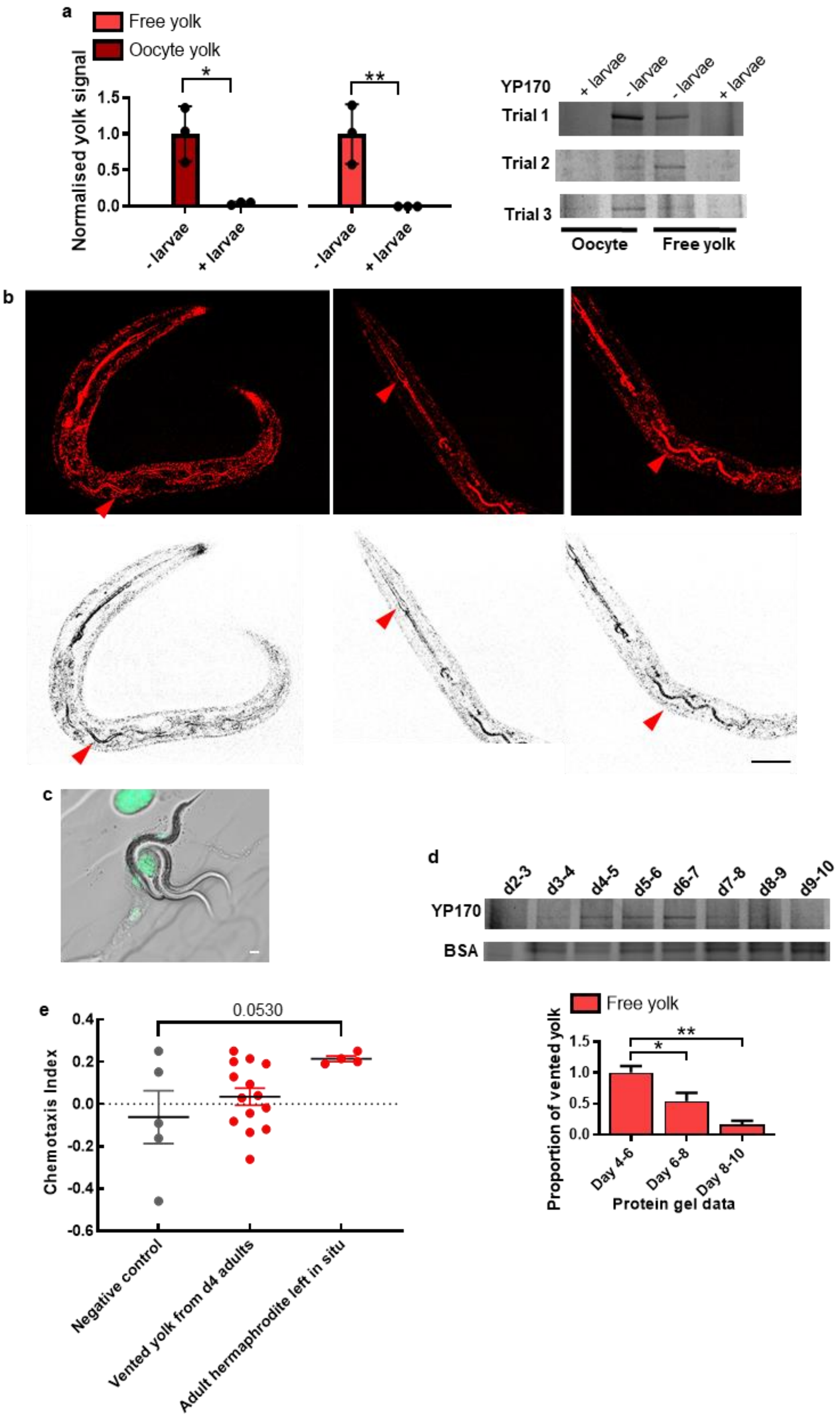


**Extended Data Fig. 2 | Larvae consume both free yolk and yolk in oocytes but are not attracted to vented yolk. a**, Larvae consume vitellogenin in both free yolk and oocyte fractions. Top: quantitated levels of YP170 from plates preconditioned with 100 venting d4 adults for 24 hr followed by addition of 200 larvae or no larvae controls for 48 hr and separation of the oocyte and free yolk fraction through centrifuging prior to protein gel electrophoresis. Data show s.d. of 3 trials. t-test, two-tailed. Bottom: protein gel image of YP170 bands from 3 trials. **b**, RCM used to highlight the dense terminal web that surrounds the intestinal lumen[18] (red arrow). Top: RCM image. Bottom: RCM image with colour inversion (dark values mapped to light and vice versa, without altering the grey distribution) to aid visualisation. **c**, Larvae on plates with no food apart from vented yolk**. d**, YP170 in free yolk protein vented by worms on different days of adulthood, and BSA standard (n=100 worms per trial). Bottom: Relative levels quantitated. Data normalised to total yolk on days 4-6. Mean ± s.e.m of 3 trials displayed. **e**, Larvae are not attracted to vented yolk (but are attracted to adult hermaphrodites). Data show calculated chemotaxis index of individual trials (each dot represents one trial) with mean ± s.e.m. displayed. One-way ANOVA. $P$ *<0.05, $P$ **<0.01.

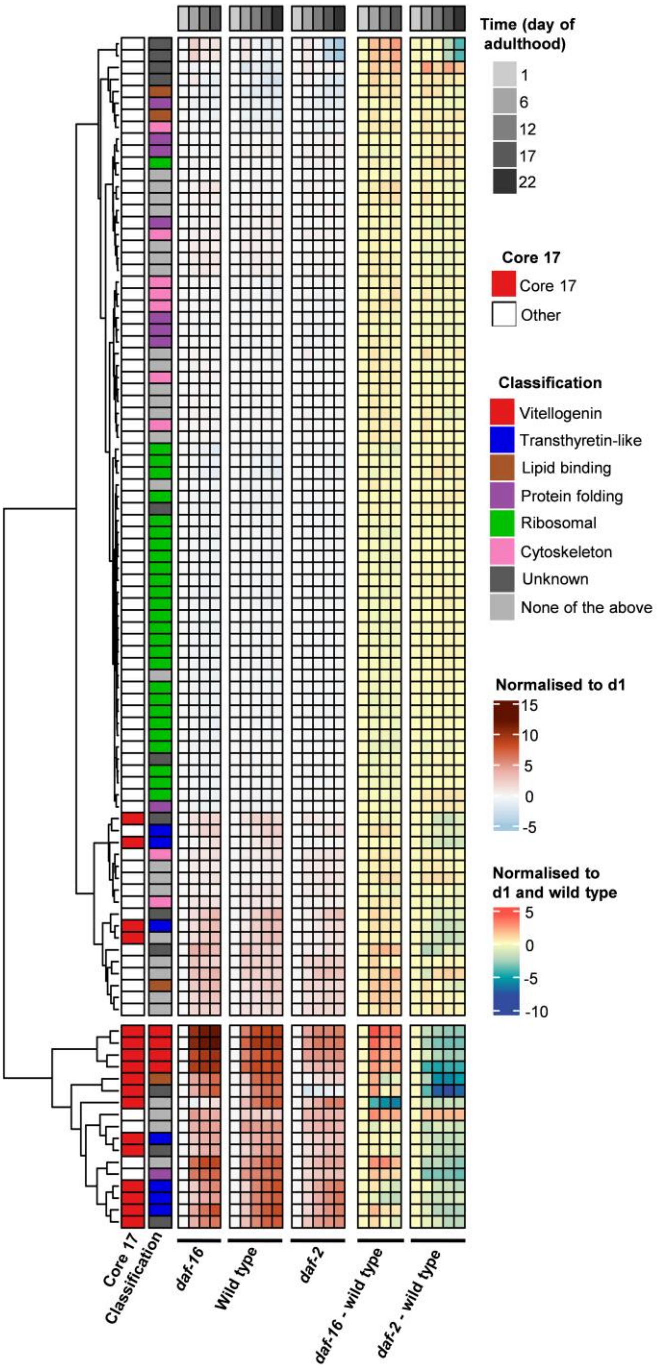


**Extended Data Fig. 3 | IIS-regulated proteins in the d4 adult-specific secretome.** Adult-specific secretome compared to published data (Walther et al., 2015) for age related changes in wild-type, *daf-2(e1370)* and *daf-16(mu86)* mutants.

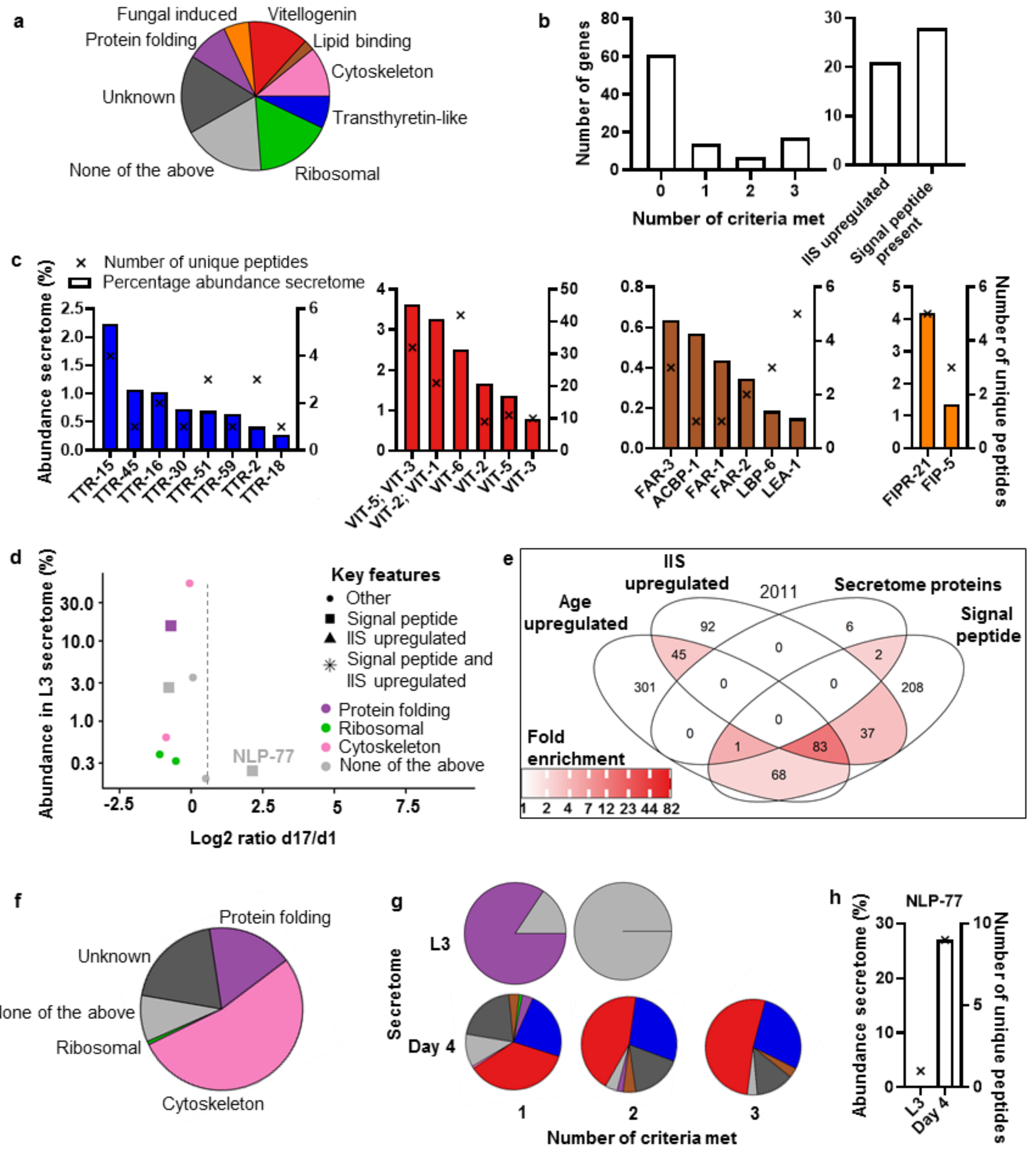


**Extended Data Fig. 4 | d4 adult specific proteome and L3 secretome comparison. a**, All proteins detected in the adult-specific d4 secretome. Manual categorisation. **b**, Number of proteins in adult specific secretome meeting only 1, only 2 or all 3 criteria of being age upregulated, IIS upregulated or possessing an N-terminal signal peptide (c.f. Extended Data Table 1), and total proteins that meet each requirement. **c**, Proteins of interest in the adult specific d4 secretome. Percentage abundance and unique peptides of all vitellogenins, transthyretins, fatty acid binding proteins and fungal induced proteins present shown. Fungal-induced proteins were not detected in the internal proteome in a previous study (Walther et al., 2015). The absence of *vit-4* unique peptides is likely an artefact resultant from the high sequence similarity between the vitellogenins: *vit-3* and *-4* have more than 99% sequence similarity to each other (Spieth and Blumenthal, 1985). A few proteotypic peptides were detected for *vit-4*. **d**, Few proteins present in the L3 secretome. Protein abundance against internal proteome age associated upregulation displayed by comparison to published data (Walther et al., 2015). **e**, No significant enrichment of proteins that are IIS upregulated and limited enrichment of age-upregulated proteins in the L3 secretome. Enrichment analysis is relative to all proteins. Significant differences in the distributions were detected using a SuperExactTest (Wang et al., 2015). **f**, The L3 secretome primarily consists of cytoskeletal and protein folding associated proteins. Percentage abundance displayed. Manual classification. **g**, Percentage abundance categorisation of proteins in L3 secretome and adult specific secretome that meet either only 1, only 2 or all 3 criteria of being age upregulated, IIS upregulated or possessing a predicted N-terminal signal peptide. No proteins in the L3 secretome meet all 3 criteria. **h**, A highly abundant neuropeptide, NLP-77, with unknown function found in the d4 secretome and also found in smaller quantities in the L3 secretome, and so not classified as part of the adult specific d4 secretome.

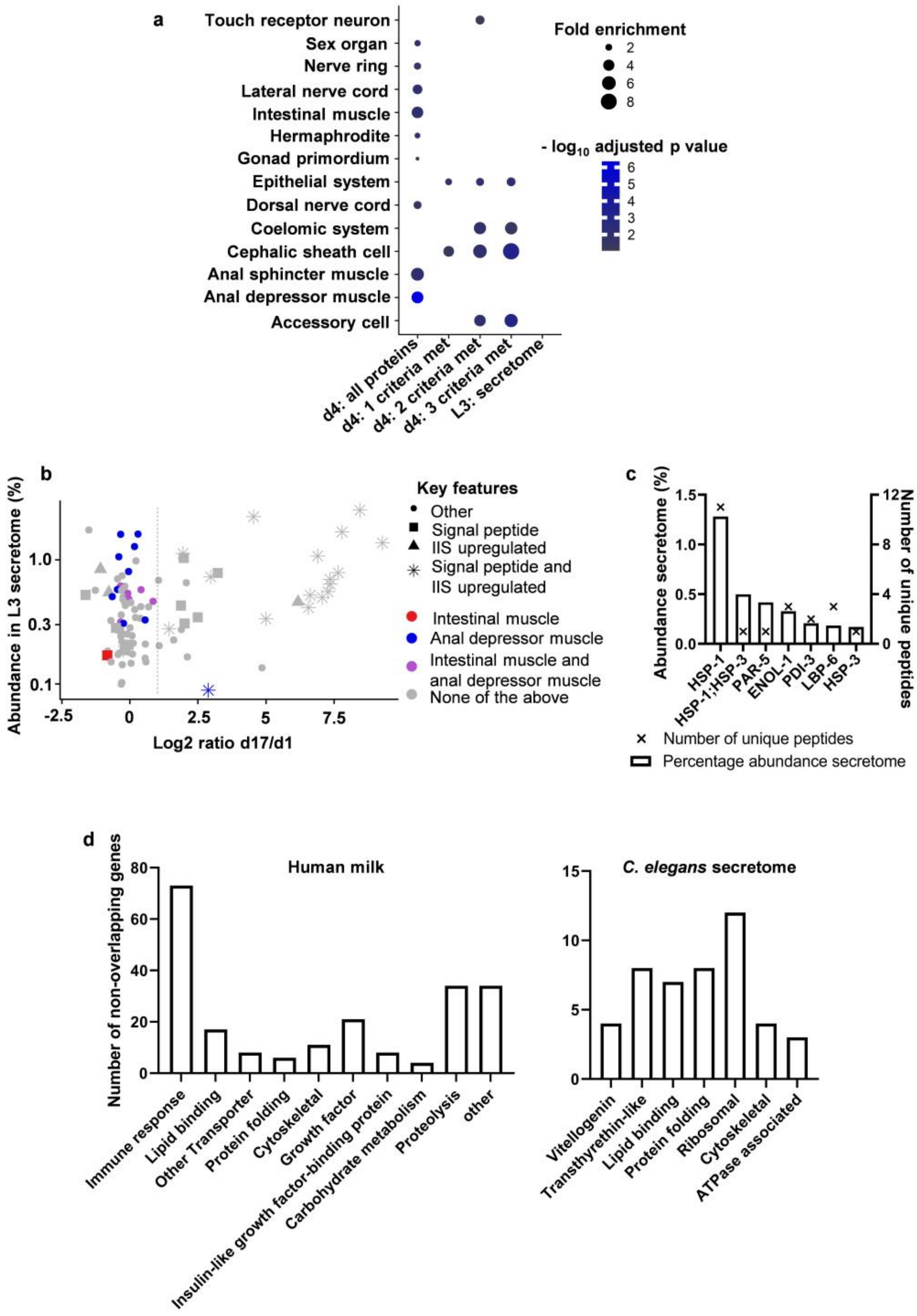


**Extended Data Fig. 5 | d4 adult specific secretome tissue enrichment analysis and comparison to the human milk proteome. a**, Tissue enrichment analysis of L3 and adult specific day 4 secretome. All proteins as well as those that met only 1, only 2 or all 3 criteria of being age upregulated, IIS upregulated or possessing a predicted N-terminal signal peptide are displayed. No enrichment was detected for the L3 secretome. Analysis was performed using the tissue enrichment analysis tool from WormBase (Angeles-Albores et al., 2016). **b**, Most anal depressor muscle-related proteins are not IIS upregulated, age upregulated or N-terminal signal peptide-containing. **c**, Conserved proteins in human breast milk and the *C. elegans* day 4 adult specific secretome. Cross-species gene set analysis performed using the XGSA package in R

(Djordjevic et al., 2016) accounting for homology mapping between *C. elegans* and humans. Human milk proteome from previously published data (D'Alessandro et al., 2010). Overlap $P$=0.00146. HSP proteins are frequently present in protein profiles, and the significant $P$ value could be due to their presence. Human homologues include: HSPA8 (heat shock protein family A (Hsp70) member 8); YWHAB (tyrosine 3-monooxygenase/tryptophan 5-monooxygenase activation protein beta); ENO1 (enolase 1); ENO2 (enolase 2); and ENO3 (enolase 3); PDIA3 (protein disulphide isomerase family A member 3); FABP7 (fatty acid binding protein 7); and HSPA5 (heat shock protein family A (Hsp70) member 5). **d**, Interpro term enrichment analysis relative to the full worm genome for the day 4 adult-specific secretome and comparison to human milk enrichment relative to human genome. Manual categorisation of groups with significant enrichment (cut-off of FDR $P$<0.05) is displayed to account for redundancy. Number of non-overlapping genes in each category is plotted. For full list of Interpro terms and Gene Ontology Biological Process terms see Supplementary File 2. Enrichment analysis was performed using Database for Annotation, Visualization and Integrated Discovery (DAVID) (Huang et al., 2009).

| | Classification | Percent intensity in secretome | Protein | Predicted function | Age increase (Log fold change over 1) | IIS upregulated | Signal peptide | Criteria met |
|---|---|---|---|---|---|---|---|---|
| P18948 | Vitellogenin | 2.51 | VIT-6 | Vitellogenin-6 | 8.45 | Yes | Yes | 3 |
| Q22288 | Transthyretin-like | 2.23 | TTR-15 | Transthyretin-like protein 15 | 4.54 | Yes | Yes | 3 |
| P05690 | Vitellogenin | 1.67 | VIT-2 | Vitellogenin-2 | 7.79 | Yes | Yes | 3 |
| P06125 | Vitellogenin | 1.37 | VIT-5 | Vitellogenin-5 | 9.28 | Yes | Yes | 3 |
| P90889 | Unknown | 1.12 | CELE_F55H12.4 | Uncharacterized protein | 1.95 | Yes | Yes | 3 |
| Q2EEM8 | Transthyretin-like | 1.07 | TTR-45 | Transthyretin-related family domain | 6.90 | Yes | Yes | 3 |
| P55955 | Transthyretin-like | 1.03 | TTR-16 | Transthyretin-like protein 16 | 1.98 | | Yes | 2 |
| G5EET8 | Unknown | 0.84 | PUD-1.2 | PUD1_2 domain-containing protein | -1.08 | Yes | | 1 |
| Q9N4J2 | Vitellogenin | 0.79 | VIT-3 | Vitellogenin-3 | 7.65 | Yes | Yes | 3 |
| Q9NA39 | Unknown | 0.78 | CCG-1 | Conserved cysteine/glycine domain protein | 3.23 | | Yes | 2 |
| Q22341 | Transthyretin-like | 0.73 | TTR-30 | Transthyretin-related family domain | 2.97 | Yes | Yes | 3 |
| O62289 | Transthyretin-like | 0.70 | TTR-51 | Transthyretin-related family domain | 7.35 | Yes | Yes | 3 |
| P52015 | None of the above | 0.69 | CYN-7 | Peptidyl-prolyl cis-trans isomerase 7 (EC 5.2.1.8) | 1.05 | | | 1 |
| Q9XW17 | None of the above | 0.65 | CAR-1 | Cytokinesis, apoptosis, RNA-associated | 2.07 | | | 1 |
| Q19478 | Lipid binding | 0.64 | FAR-3 | Fatty acid/retinol binding protein | 7.36 | Yes | Yes | 3 |
| G5EDZ9 | None of the above | 0.55 | CPI-1 | Cystatin (Cysele1) | 7.27 | Yes | Yes | 3 |
| G5EBF3 | Unknown | 0.55 | PUD-2.1 | Protein up-regulated in *daf-2(gf)* | -0.77 | Yes | | 1 |
| O16462 | Unknown | 0.52 | GRD-5 | Ground-like domain-containing protein | -1.62 | | Yes | 1 |
| Q23683 | Unknown | 0.52 | CELE_ZK970.7 | DUF148 domain-containing protein | 6.62 | Yes | Yes | 3 |
| Q19063 | Unknown | 0.50 | CELE_E04F6.8 | Uncharacterized protein | 7.07 | Yes | Yes | 3 |
| Q20363 | Protein folding | 0.46 | SIP-1 | Stress-induced protein 1 | 6.18 | Yes | | 2 |
| O18089 | Unknown | 0.43 | CELE_T13F3.6 | DUF19 domain-containing protein | 1.88 | | Yes | 2 |
| P34500 | Transthyretin-like | 0.41 | TTR-2 | Transthyretin-like protein 2 | 6.56 | Yes | Yes | 3 |
| P34383 | Lipid binding | 0.34 | FAR-2 | Fatty-acid and retinol-binding protein 2 | 2.49 | | Yes | 2 |
| Q19064 | Unknown | 0.33 | CELE_E04F6.9 | Uncharacterized protein | 5.00 | Yes | Yes | 3 |
| O45599 | None of the above | 0.31 | CBD-1 | Chitin-binding domain protein | 2.01 | | Yes | 2 |
| O01504 | Ribosomal | 0.28 | C37A2.7 | 60S acidic ribosomal protein P2 | -0.51 | | Yes | 1 |
| Q17473 | Transthyretin-like | 0.28 | TTR-18 | Transthyretin-related family domain | 1.44 | Yes | Yes | 3 |
| G5EEA8 | None of the above | 0.27 | NEX-1 | Annexin | 1.89 | | | 1 |
| Q21763 | None of the above | 0.23 | CELE_R05H5.3 | Thioredoxin domain-containing protein | 1.60 | | | 1 |
| G5ED07 | Protein folding | 0.21 | PDI-3 | Protein disulfide-isomerase (EC 5.3.4.1) | -0.05 | | Yes | 1 |
| Q07750 | Cytoskeleton | 0.20 | UNC-60 | Actin-depolymerizing factor 1, isoforms a/b | 1.07 | | | 1 |
| Q17967 | Protein folding | 0.19 | PDI-1 | Protein disulfide-isomerase 1 (PDI 1) (EC 5.3.4.1) | -0.17 | | Yes | 1 |
| Q20724 | Unknown | 0.19 | CELE_F53F4.13 | Uncharacterized protein | -0.24 | Yes | Yes | 2 |
| P27420 | Protein folding | 0.17 | HSP-3 | Heat shock 70 kDa protein C | -0.83 | | Yes | 1 |
| P91306 | None of the above | 0.16 | CEY-2 | CSD_1 domain-containing protein | 2.11 | | | 1 |
| O17687 | None of the above | 0.13 | NASP-2 | SHNi-TPR domain-containing protein | 4.85 | | | 1 |
| Q21265 | None of the above | 0.09 | TAG-225 | Putative metalloproteinase inhibitor (TIMP-like protein) | 2.88 | Yes | Yes | 3 |

**Classification**

- Vitellogenin
- Transthyretin-like
- Lipid binding
- Protein folding
- Ribosomal
- Cytoskeleton
- Unknown
- None of the above

**Extended Table 1 | d4 adult specific secretome proteins that are either IIS upregulated, age upregulated or have a predicted N-terminal signal peptide.** Significant differences in the distributions were detected using a Kolmogorov-Smirnov test.
